# Genome sequence of *Talaromyces trachyspermus*, a biocontrol fungus isolated from broomrape

**DOI:** 10.1101/2025.07.07.663553

**Authors:** Roghayeh Hemmati, Aria Dolatabadian, Sobhan Saeedi, Jacqueline Batley

## Abstract

Applying antimicrobial compounds derived from microorganisms for plant disease management is one of the objectives of sustainable agriculture. The genus *Talaromyces* is known for its species’ ability to produce a diverse group of antimicrobial compounds. For example, *T. trachyspermus* has been reported to produce secondary metabolites, cell wall-degrading enzymes, and plant growth-promoting factors. Identification of novel promising metabolites and enzymes from *T. trachyspermus* is still in its infancy. Also, there is a lack of information about the genomic resources for its secondary metabolites and hydrolytic enzymes. Therefore, this study aimed to analyse the genome of a biocontrol isolate of this species to investigate its biocontrol mechanisms at the genomic level, focusing on secondary metabolites and cell wall degrading enzymes. The whole genome of *T. trachyspermus* isolate IRAN 3054C, obtained from necrotic *Orobanch ramosa* stems in Iran with biocontrol ability, was sequenced using the Illumina platform. We performed both *de novo* and resequencing analyses of the genome, obtaining a 31.3 Mb assembly. The abundance of protein groups associated with biocontrol activities was assessed in the studied genome. Fungismash was used to detect and annotate secondary metabolites. The analysis revealed the presence of several secondary metabolite biosynthesis gene clusters (BGCs), with a high frequency of polyketide synthases (T1PKS) and nonribosomal peptide synthetases (NRPS), which are known to produce bioactive compounds with antimicrobial properties. Among the identified secondary metabolites, Fusarin, YWA1, Dimethylcoprogen, and Squalestatin S1 exhibited the highest similarity to known compounds. Furthermore, sequences similar to Phyllostictine A/B and Cornexistin indicate potential herbicidal properties. The genome also had domains for enzymes involved in phosphate solubilisation, siderophore production, and fungal cell wall degradation, which are essential for biocontrol and plant growth promotion. Our findings highlight the genomic richness of *T. trachyspermus* IRAN 3054C for biocontrol. Further metabolomics studies are needed to validate the actual production of these secondary metabolites and explore their functional roles in biocontrol.

## 1- Introduction

*Talaromyces* belonging to Ascomycota, Eurotiomycetes, Trichocomaceae, is a teleomorphic form for some *Penicillium* species. Based on multi-locus phylogenetic analysis, the genus is classified into seven sections (Yilmaz et al., 2014). *T. trachyspermus* (Shear) Stolk & Samson, 1972, is a species within the *Talaromyces* section *Trachyspermi*, distinguished by slow-growing, floccose white to yellow mycelium, mono-verticillate conidiophores, smooth ellipsoidal conidia, and abundant globose to sub-globose cleistothecial ascomata with ellipsoidal ascospores (Stolk & Samson, 1972). *Talaromyces* species are distributed in nature and different environments. They have been isolated from various substrates such as soil, food, and plant materials. Some species of this genus produce anti-cancer, anti-bacterial, and antifungal compounds; others are important producers of lignocellulolytic enzymes and natural pigments (Fu et al., 2024). Some plant-associated species have been reported as endophytes (Yilmaz et al., 2016; Quan et al., 2024), whereas some are plant-pathogenic fungi (Sun et al., 2020). Some species are mycotoxin producers, and others cause food spoilage (Yilmaz *et al*., 2014).

Several reports show the antagonistic effects of Talaromyces species against plant pathogenic fungi and /or their plant growth-promoting effects. (Fu et al., 2024; Farhat et al., 2023; Zhang et al., 2024; Manoch and Dethoup, 2011; Nagtzaam, 1998; Goh et al., 2020; Fahat *et al*., 2021; Sahu *et al*., 2019; Zhao *et al*., 2022; Abdel-Rahim and Abo-Elyousr, 2018; Fahima and Henis, 1995). Among those species, *T. trachyspermus* is an important *Talaromyces* species of antagonistic fungi. There are reports of its biocontrol effects against plant pathogenic fungi, including *Alternaria brassicicola, Colletotrichum capsici, Pythium aphanidermatum, Rhizoctonia solani, Sclerotium rolfsii* and *Scleortinia sclerotiorum, Alternaria alternata,Alternaria arborescens, Botryosphaeria dothidea* and *Colletotrichum gloeosporioides* (Dethoup *et al*., 2015; Sahu *et al*., 2019; Zhao *et al*., 2020). One isolate of *T. trachyspermus*, isolated from the necrotic tissue of broomrape stems (*Orobanche* spp.) in Iran, caused a significant reduction in the number of tubercles, indicating its potential as a biocontrol fungus against this parasitic plant (Hemmati and Gholizadeh, 2019). Several other fungal species, specifically within the *Fusarium* genus, have been reported as mycoherbicidal fungi against *O. ramosa* (Abouzeid and El-Tarabily, 2010; Ghannam *et al*., 2007; Abouzeid *et al*., 2004; Boari and Vurro, 2004). *T. trachyspermus* isolated from the medicinal plant *Withania somnifera* (winter cherry) showed high production of hydrolytic enzymes, protease, chitinase, amylase, cellulase, and pectinase, which are required for biocontrol activities. In the meantime, this isolate produced high levels of indole acetic acid, siderophore synthesis, and phosphate solubilisation activities important for plant growth promotion (Sahu *et al*., 2019). This species has been reported as a successful producer of Spiculisporic acid (SA) as a fatty acid-type biosurfactant useful in the cosmetics industry (Moriwaki-Takano *et al*., 2021).

*T. trachyspermus* has gained attention recently due to its production of various bioactive secondary metabolites with antimicrobial properties (Farhat *et al*., 2022; Zhai *et al*., 2016). Secondary metabolites (SMs) are a group of low molecular weight natural products that are not vital for their producers but can increase their fitness in different habitats. Consequently, they can increase their survival in competitive environments (Conrado et al., 2022). Fungal secondary metabolites are divided into four classes: polyketides, non-ribosomal peptides, terpenoids and shikimic acid derived compounds (Pusztahelyi et al., 2015). Polyketides are a large class of secondary metabolites produced by bacteria, fungi, plants, and a few animal groups (Roy, 2019). They are used as immunosuppressants, antibiotics, cholesterol-lowering agents, and anticancer agents (Roy, 2019). Nonribosomal peptide synthetases (NRPS) are enzymes synthesizing a variety of peptides. Many of these peptides have biological and biotechnological potential. Some have antimicrobial activity, such as penicillin and vancomycin, and anticancer features, such as bleomycin (Iacovelli et al., 2021). Some fungal secondary metabolites are toxic to plants. Phytotoxins are fungal secondary metabolites (FSMs) that can be classified into two groups: host-specific toxins (HST) and non-host-specific toxins (non-HST). Among them are some toxins such as T-toxin, HC-toxin, Victorins as HSTs and Tentoxin and Cercosporin as non-HSTs (Stergiopoulos et al., 2013). Several secondary metabolites from fungi have been reported as potential mycoherbicides. For example, neosolaniol monoacetate, a trichothecene isolated from cultures of a *F. compactum* strain infecting *O. ramose* tissues, was evaluated for its ability to inhibit the seed germination of parasitic plants (Andolfi et al. 2005). A number of microbial products from fungi, such as gulfosinate, tentoxin, cornexistin, moniliformin, and fusaric acid, have been successfully used to manage different weeds (Singh and Pandey, 2019). Certain FSMs enhance plant growth and stress tolerance. Auxin-like substances produced by some endophytic fungi play a role in plants’ root growth and biomass deposition (Gravel et al., 2007). Some fungal secondary metabolites have antimicrobial properties that protect plant hosts from invading pathogens. For instance, *Trichoderna* is capable of producing hundreds of anti-microbial secondary metabolites, including Trichodermin (Buller et al., 1971), trichomycin, chlorotrichomycin, gelatinomycin, antibacterial peptides (Maruyama et al., 2020) and pentaibols. These secondary metabolites can act as antibacterial agents against bacterial and fungal pathogens, including *Fusarium oxysporum* and *Phytophthora nicotiana* (Yao et al., 2023). The genes encoding enzymes for synthesis of secondary metabolites are grouped into secondary metabolites biosynthesis gene clusters (SM-BGCs). The secondary metabolites of *Talaromyces* mainly include alkaloids, peptides, lactones, polyketides, and miscellaneous structure-type compounds (Zhai et al., 2016). Genomic analysis of some of the *Talaromyces* species has provided useful information about the genetic determinants of secondary metabolite biosynthesis. *The talaromyces pinophilus* strain 1–95 genome was sequenced and revealed a high number of secondary metabolite gene clusters. They discovered 68 gene clusters for secondary metabolism consisting of type I polyketide synthase (T1PKS) genes and nonribosomal peptide synthetase (NRPS) genes. These gene clusters are important in producing a variety of bioactive compounds, including antimicrobials essential for antagonistic activity (Li et al., 2017). In another study, the endophytic fungus *Talaromyces* sp. strain DC2 had 20 biosynthetic gene clusters for secondary metabolite production. The gene list indicates that the strain can produce a broad spectrum of secondary metabolites (Quan et al., 2024). The genome of *T. albobiverticillius* Tp-2 contains eight distinct gene clusters for secondary metabolite biosynthesis. This strain showed a high genomic capacity to generate different bioactive compounds, including pigments with the potential of industrial applications (Wang et al., 2023). The production of cell wall degrading enzymes (CWDEs), including proteases, chitinases and glucanases, is one of the essential mechanisms that fungal biocontrol agents employ against plant pathogenic fungi (Inglis and Kawchuk, 2002). The different categories of carbohydrate-active enzymes CAZymes are grouped into different families, including but not limited to glycoside hydrolases (GHs), polysaccharide lyases (PLs), carbohydrate esterases (CEs) and carbohydrate-binding modules (CBMs) (Cantarel et al., 2009). All fungal chitinases described so far are members of GH family 18 (Seidl, 2008). In addition to chitin as a main structural component of fungal cell wall, the second group of fibrillar polymer in fungal cell walls are glucans (Latgé, 2007); glucanases have been placed into GH families 16, 55, 64, 81, 5, 30,27, 71 and 92 (Kubicek et al., 2011; Gruber and Seidl-Seiboth, 2012). Besides carbohydrate-active enzymes, proteases are important for deleting the fungal cell wall. It has been demonstrated that mycoparasite species of *Trichoderma* have more proteases, a higher number of chitinase coding genes, and more copies of glucanases compared with saprophytic species (Gruber and Seidl-Seiboth, 2012).

Several works have reported high potential for production of biomass degrading enzymes by different Talaromyces species. Genomic analysis of *T. pinophilus* revealed that this species has 803 genes to encode enzymes acting on carbohydrates; among them, 39 enzymes were cellulose-degrading, and 24 were starch-degrading (Li et al., 2017). A whole genome analysis of *Talaromyces* sp. strain DC2 revealed that the genome of this strain has a total of 149, 227, 65, 153, 53, and 6 genes responsible for cellulose, hemicellulose, lignin, pectin, chitin, starch, and inulin degradation, respectively (Quan et al., 2024). There are several works on biochemical identification and product optimization of some important enzymes from different species of this genus including xylanase production by *T. amestolkiae* on agro-industrial wastes (Barbieri et al., 2022), GH51 α-l-arabinofuranosidase production by *T. leycettanus* (Tu et al., 2019), industrially significant proteases including thermostable aspartic protease from *T. leycettanus* (Guo et al., 2020), a novel AA14 LPMO from *T. rugulosus* with strong oxidative activity on cellulose, xylan and xyloglucan (Chen et al., 2024) and identification of a gene encoding an extracellular β-galactosidase of *T. cellulolyticus* (Orleneva et al., 2022).

Genome-wide analysis and other bioassay and biochemical experiments such as metabolomics will help to understand the biocontrol mechanisms such as antibiosis and parasitism and the genomic basics underlying these mechanisms. For *T. trachyspermus*, only one genome assembly of 32Mb size belonging to strain 4014 is isolated from a medicinal plant (Sahu et al., 2019). There is no genome annotation and published data analysis for this whole genome sequence. Therefore, there is a lack of information regarding the genomic resources for biocontrol detreminants such as secondary metabolites, cell wall degrading enzymes and plant growth promotion factors for this species. In the current study, we performed whole genome sequencing for a biocontrol isolate of this species, IRAN 3054C, which we have isolated from broomrape in Iran (Hemmati and Gholizadeh, 2019). This isolate have shown antifungal effects, biocontrol activity against *O. ramosa*, and plant growth promotion phenotypes (Hemmati and Gholizadeh, 2019 and unpublished data). As a part of whole genome data analysis, we aimed to predict proteins and secondary metabolites, specifically carbohydrate-active enzymes (CAZYmes) and polyketides which are associated with biocontrol. The genomic analysis of this fungal isolate will help us to have a deeper and more precise understanding of its capacities as a promising biocontrol agent.

## 2- Materials and methods

### 2-1- The origin of the isolate and preparation of fresh cultures

This study used a *T. trachyspermus* isolate (IRAN 3054C) obtained from broomrape plants showing stem rot symptoms. This isolate has previously been reported as a potential biocontrol agent against broomrape due to its effects on broomrape seedlings and a significant reduction in the number of emerging shoots of this pathogenic plant in tomato pots under greenhouse conditions (Hemmati and Gholizadeh, 2019). The dehydrated form of this isolate was preserved at -20°C as dried pellets of potato dextrose agar (PDA) cultures of the fungus. A small pellet was placed on a Petri dish containing PDA and incubated in darkness at room temperature (24 ± 2 °C). After ten days, the fresh cultures were ready for DNA extraction.

### 2-2- DNA extraction, qualification and quantification

The isolate was cultured on PDA, and seven-day-old colonies were stored at -70°C for 24 hours before DNA extraction. Then, the frozen mycelia were harvested from the culture media by scraping the hyphal layer from the surface of the agar medium and ground by using sterilised porcelain mortar and pestel. The DNA was extracted using the CTAB method (Moller *et al*., 1992). The total genomic DNA was assessed for quality using 1% agarose gel electrophoresis and was quantified using a Qubit.

### 2-3- Library preparation and whole genome sequencing

The minimum volume of 10 ul of each DNA sample, with a minimum concentration of 10 ng.ul^-1^, was sent to AGRF (Australian Genome Research Facility) for library preparation and whole-genome sequencing. Sequencing was performed on the Illumina platform with two paired-end reads (150 bp), resulting in a coverage of 20.

### 2-4- Whole-genome data analysis

#### 2-4-1- Resequencing analysis

A FASTA format of a reference genome for *T. trachyspermus* (BUMICRO_TalaroTrachy_1.1, strain 4014) was downloaded from NCBI. The related FASTA file was uploaded to “use.Galaxy.org” an annotated GFF3 file was created using Augustus. The resulting file was imported into the CLC Genomics Workbench 20.0 (QIAGEN, Denmark). For mapping to the reference genome, the genome of our biocontrol isolate was aligned with the reference genome of *T. trachyspermus* (strain 4014) using CLC Workbench. Variant calling was performed to identify single nucleotide polymorphisms (SNPs) and small insertions/deletions (indels) between the biocontrol isolate and the reference genome.

#### 2-4-2- *de novo* assembly

The raw sequencing data underwent quality control and preprocessing steps in the CLC workbench. This involved assessing sequencing quality and trimming low-quality bases. A *de novo* assembly was performed on the resulting reads in the CLC workbench to obtain some genomic features of our isolate. This Whole Genome Shotgun project has been deposited at DDBJ/ENA/GenBank under the accession JBNAEF000000000. The version described in this paper is version JBNAEF010000000. The associated BioProject and BioSample accessions are PRJNA1243591 and SAMN47626500, respectively. The percentage of repetitive sequences in the genome was identified using the Red repeat masking pipeline (Red, version 2018.09.10, Girgis, 2015) available on the Galaxy web-based analysis platform. As a gene prediction tool on the Galaxy platform, Augustus was used for protein-coding gene prediction.

Further quality assessments were performed by using the BUSCO pipeline in the Galaxy web-based analysis platform with ‘eurotiales_odb10’ and used as reference datasets (https://busco.ezlab.org/). The BUSCO pipeline was also used to predict and annotate protein-coding genes using the default e-value threshold of <1e-03. The BUSCO annotation pipeline utilized metaeuk (v5.34c21f2) to annotate the fungal assemblies (Levy Karin et al., 2020).

EggNOG (v2.1.8) in the online GALAXY platform was used to identify and annotate Orthologous Groups (OGs) using the default e-value threshold of 10^−3^. This analysis provided several forms of functional annotation, including Gene Ontology terms, KEGG pathways, SMART/Pfam domains, and Clusters of Orthologous Groups of proteins (COGs) (Tatusov *et al*., 2000). To analyse the Gene Ontology (GO) terms for all the predicted Pfam domains, we mapped them against the data in STRING (<u>STRING: functional protein association networks (string-db.org)</u>). The web-based program Categorizer was used to classify the GO terms for all the identified domains (Hu *et al*., 2008). Genes belonging to 13 protein groups were selected based on their reported involvement in biocontrol-associated functions, such as parasitic activities, and their abundance over the genome was recorded for our studied isolate.

Fungismash (https://fungismash.secondarymetabolites.org/) was used to detect and functionally annotate secondary metabolites encoding gene clusters (SM-BGSs) using the strict detection strictness setting (Blin *et al*., 2021).

## 3- Results

### 3-1- The results of variant callings for re-sequencing analysis

After mapping the reads to the *T. trachyspermus* (strain 4014) as the reference genome, the total number of variants was 312,928, including 96,634 insertions, 11,121 deletions, 190,295 single nucleotide variants (SNVs), 12,671 multi nucleotide variants (MNVs), and 2,207 replacements. Among the variants, 28,571 (9.13 %) were associated with amino acid changes in the studied isolate, including 132 replacements, 10,893 insertions, 15,666 SNV, 1,243 MNV and 639 deletions.

### 3-2- Genome features following *de novo* assembly

A 31.31 Mb genome was generated with 20-fold coverage. The size was smaller than the estimated genome size of 32 Mb for *T. trachyspermus* strain 4014, which serves as the reference genome for this species. Five hundred fifty-two contigs covered the genome. The N50 and N75 sizes of the contigs were, respectively, 385.98Kb and 176.13Kb. The GC content was 47.3 %. The BUSCO completeness score for the studied genome was 92.26%, with a total of 4,191 BUSCO genes identified (Table 1).

**Table 1:** Summary of *Talaromyces trachyspermus* strain IRAN 3054C genome sequencing and assembly results.

|  |  |
| --- | --- |
| Total sequenced bases | 1,594,985,700 |
| Mean read length | 150 |
| Number of reads | 10,633,238 |
| Contigs | 552 |
| Largest contig (bp) | 1,321,046 |
| Total length | 31,309,464 |
| GC (%) | 47.3 |
| N50 | 385,981 |
| N75 | 176,132 |
| Protein-coding genes (from Augustus) | 9562 |
| Repeats content (%) | 59 |
| Busco annotation results (%) |  |
| Complete | 92.26 |
| Complete and single copy | 91.26 |
| Complete and duplicate | 1 |
| Fragmented | 0.85 |
| Missing | 6.89 |

### 3-3- Clusters of Orthologous Groups

Functional annotation analysis (EggNOG) detected 8,503 orthologous groups (OGs) in the *Talaromyces* strain; among them, 95.2% of OGs were assigned a COG annotation (Clusters of Orthologous Groups of proteins). As depicted in Figure 1, the COG functional categories with the highest mean relative abundances included ‘function unknown’ (24.56%), ‘Carbohydrate transport and metabolism’ (7.09%), ‘Secondary metabolite biosynthesis, transport, and catabolism’ (6.71%) and ‘amino acid transport and metabolism’ (6.39%).

**Figure 1:**
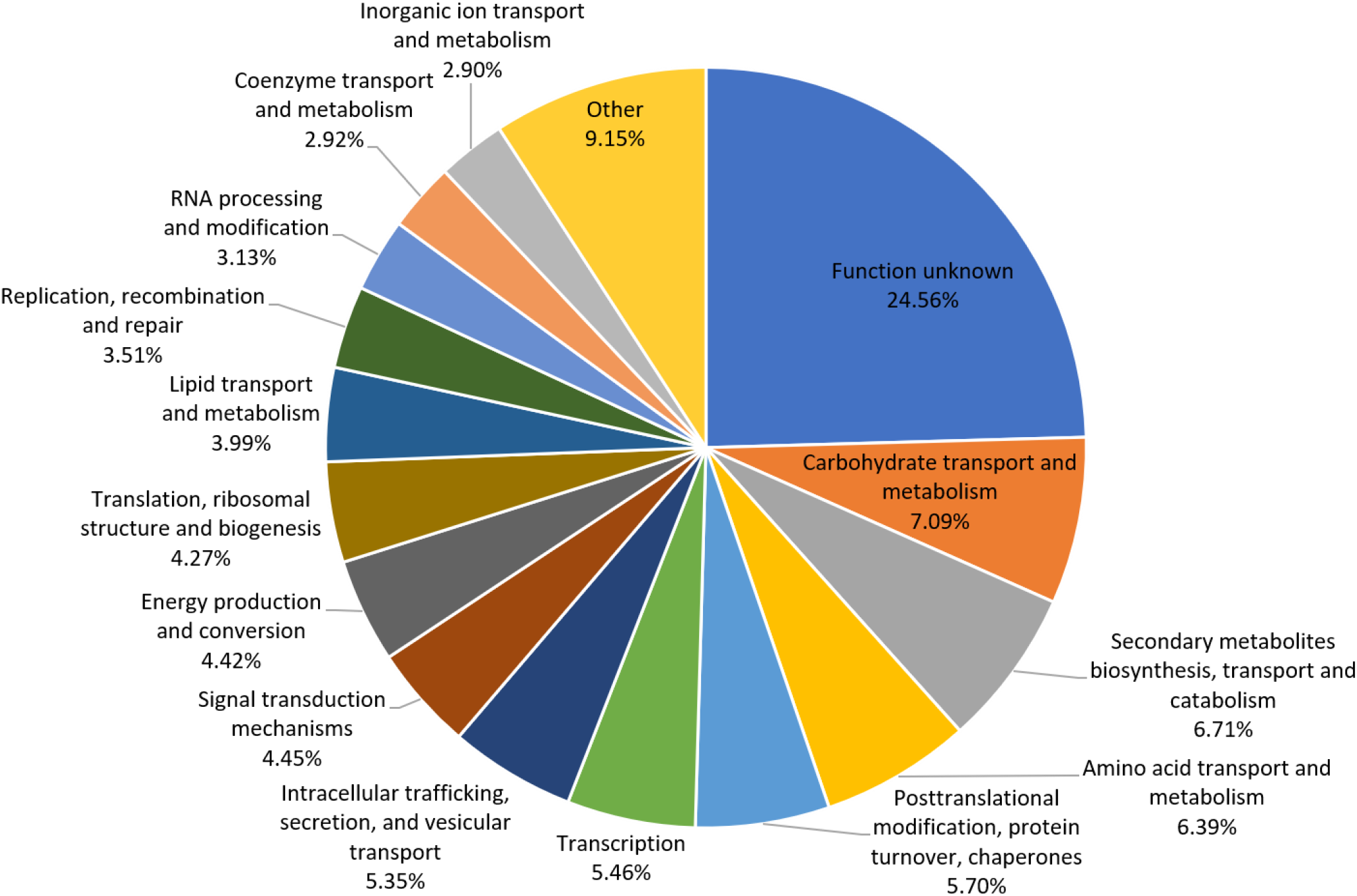
Clusters of Orthologous Groups (COG) functional categories and their abundances (%) in the *Talaromyces trachyspermus* Iran IRAN 3054C genome. For display limitations, nine of the COG functional categories with a mean relative abundance of less than 2% were grouped into the category of ‘Other’.

By mapping identified GO terms for the studied isolate to 127 of the GO_slime ancestor terms by a single count, available at CateGOrizer (Hu et al., 2008), 182 GO terms were classified into biological process, 127 into metabolism, 88 into cellular components and 31 into molecular function. Overall, we had 301 unique terms belonging to at least one of the 45 GO_slime classes (Figure 2).

**Figure 2:**
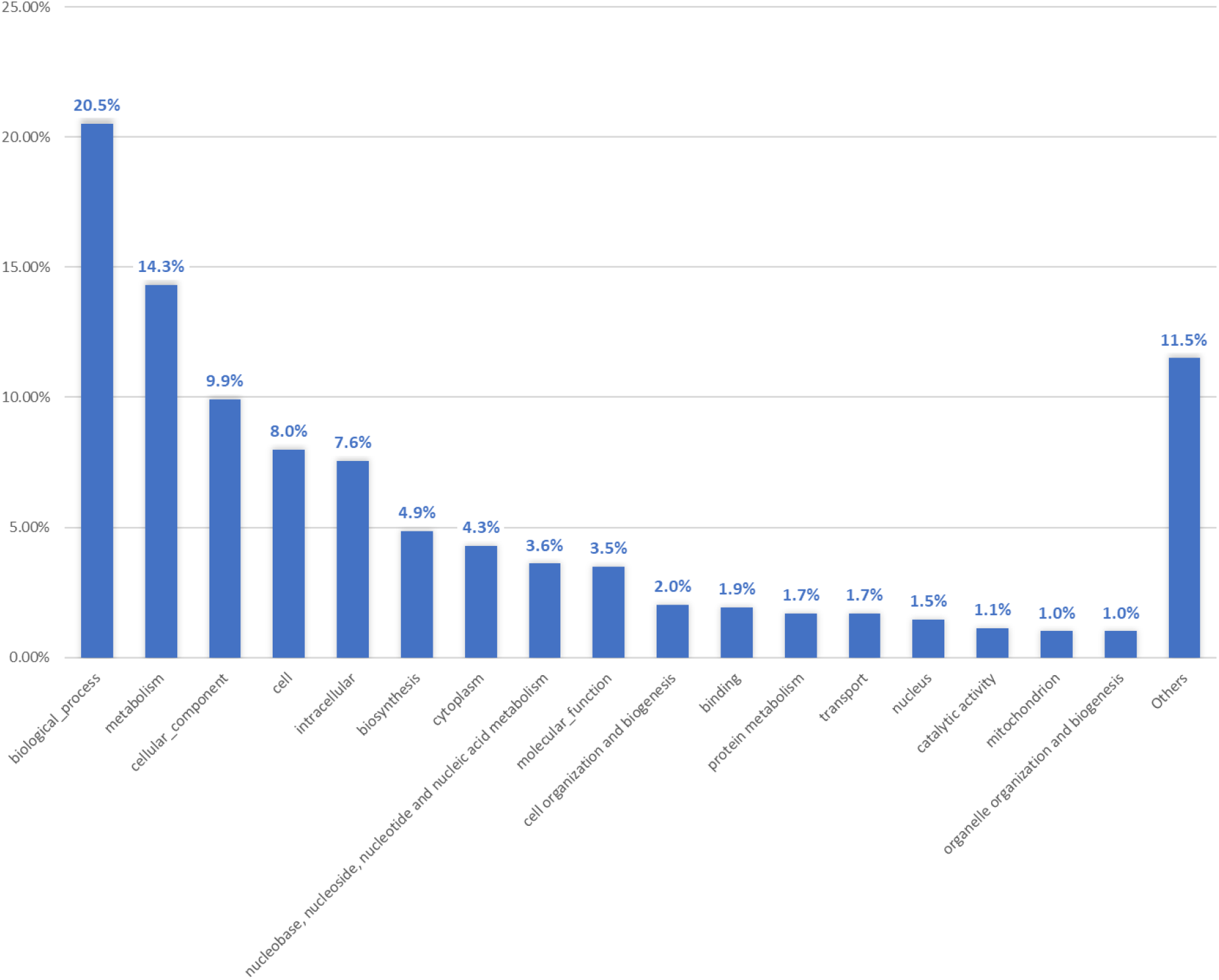
Go terms classification count resulted by mapping identified GO terms to GO_slime ancestor terms, available at CateGOrizer.

Among the Pfam domains, we identified several domains for producing carbohydrate-active enzymes (CAZymes) and fungal cell wall degrading enzymes (FCWDEs). Peptidases had the most domains in our studied genome (count number: 93). The second and third most frequent domains with fungal cell wall degrading activity or parasitism belonged to chitinases (count number: 31), proteases (30), and lipases (30). Glucanases, the other group of FCWDEs, had 13-count domains in the genome (Table 2).

**Table 2:** Pfam domains associated with carbohydrate-active enzymes, fungal cell wall degrading enzymes, and plant growth-promoting factors in the *Talaromyces trachyspermus* genome extracted from EggNOG output.

| Pfam Domain | Function / Process | The count number of domain |
| --- | --- | --- |
| β-glucosidases | Synthesis and degradation of complex carbohydrates (CAZyme) | 7 |
| Glucanases | CAZyme<sup>1</sup> / FCWDEs<sup>2</sup> | 13 |
| Galactosidases | CAZyme | 4 |
| Chitinases (GH18) | CAZyme/ FCWDEs | 31 |
| Chitosanases | CAZyme / FCWDEs | 0 |
| Mannosidases | CAZyme | 10 |
| α-L-Arabinofuranosidase | CAZyme | 1 |
| β-L-Arabinofuranosidase | CAZyme | 3 |
| Glycoside hydrolase: Xylanase | CAZyme | 8 |
| N-acetyl-β-D-glucosaminidase | CAZyme | 1 |
| Cellolytic enzymes | CAZyme | 3 |
| Pectate lyase | CAZyme | 6 |
| Peptidase | Parasitism / FCWDEs | 93 |
| Protease | Parasitism / FCWDEs | 30 |
| Lipase | Parasitism | 30 |
| Siderophore synthase | PGP | 5 |
| Indole acetic acid | PGP | 0 |
| Acid phosphatases | PGP | 15 |
| Alkaline phosphatases | PGP | 2 |
| Phytase | PGP | 1 |

1: Carbohydrate Active Enzymes; 2: Fungal Cell Wall Degrading Enzymes; 3: Plant growth promotion.

### 3-4- Secondary metabolite gene clusters

FungiSMASH analysis identified 18 different types of secondary metabolites biosynthesis gene cluster (SM-BGC) classes in the genome of *T. trachyspermus* (Table 3, Table 4), including T1PKS, terpenes, NRPS, NRPS-like, β-lactones, fungal-RiPP-like, and hybrid BGCs, including T1PKS/NRPS and T1PKS/NRPS-like. Forty-six BGCs were detected in this studied genome (Table 3). Four BGCs were identified to share 100% similarity to known BGCs that encode for fusarin, dimethylcoprogen, YWA1, and choline biosynthesis. Four BGCs were identified to share at least 40% but less than 100% similarity to known BGCs that are deposited in the MIBiG repository, including squalstatin S1, nidulanin A, phomoidride and phyllostictine A/phyllostictine B (https://mibig.secondarymetabolites.org). A further 10 BGCs were identified to share 7% to 33 % similarity to known BGCs (Table 4). Most of the BGCs could not be identified in our studied species. The most abundant SM-BGC in the studied genome belonged to Type 1 polyketide synthases (T1PKSs).

**Table 3:**
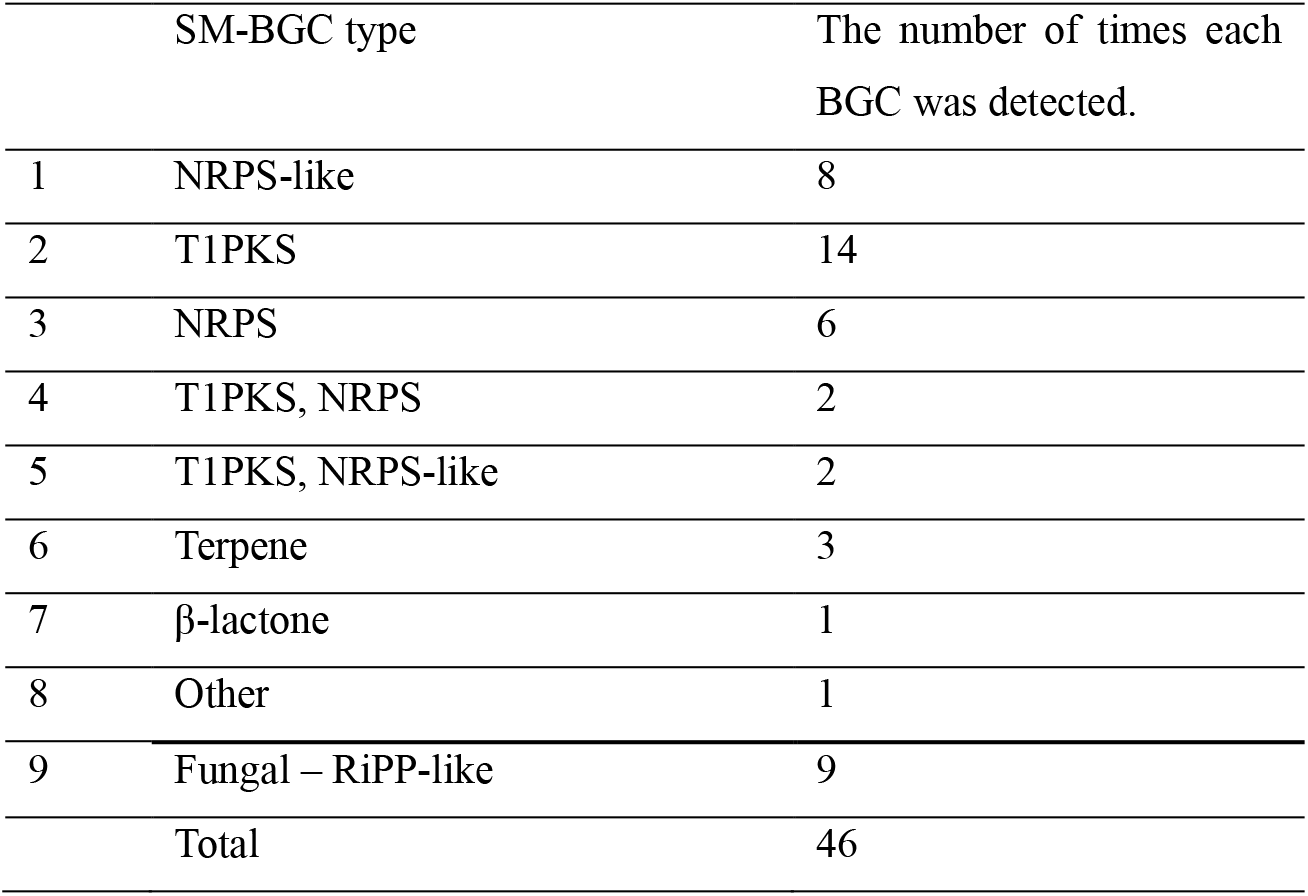
The number of times that each type of SM-BGC was detected in the genome of *Talaromyces trachyspermus* strain IRAN 3054C by FungiSMASH analysis.

**Table 4:** The secondary metabolites that were detected in the genome of *Talaromyces trachyspermus* by fungiiSMASH analysis.

| BGC type | Length (nt) | Most similar compound | MIBiG accession | Similarity (%) |
| --- | --- | --- | --- | --- |
| Terpen | 21484 | Squalestatin S1 | BGC0001839 | 60 |
| NRP | 49,455 | Nidulanin A | BGC0001699 | 50 |
| Polyketide | 48,197 | Phomoidride | BGC0001913 | 46 |
| NRP+Polyketide | 46,189 | Fusarin | BGC0000064 | 100 |
| NRP+Polyketide | 50,496 | Phyllostictine A/phylllostictine B | BGC0001741 | 40 |
| NRP+Polyketide | 52,406 | Phyllostictine A/phylllostictine B | BGC0001741 | 30 |
| Polyketide | 45991 | Ywa1 | BGC0002175 | 100 |
| NRP | 26,233 | Dimethylcoprogen | BGC0001249 | 100 |
| NRP | 43,831 | Choline | BGC0002276 | 100 |
| Polyketide | 43,912 | Chromane | BGC0001907 | 33 |
| Polyketide | 47,709 | Depudecin | BGC0000046 | 33 |
| NRP | 41,667 | Metachelin | BGC0002710 | 25 |
| NRP+Alkaloid+Polyketide:Iterative<br>type I polyketide | 51,435 | Ucs1025a | BGC0001449 | 23 |
| Polyketide | 46,877 | Scytalidin | BGC0002223 | 18 |
| Polyketide | 44,195 | Aurofusarin | BGC0002709 | 18 |
| Polyketide | 37,812 | Cornexistin | BGC0001436 | 14 |
| Polyketide | 44,195 | Elsinochrome C | BGC0001865 | 12 |
| Terpene | 40,081 | Demethoxyviridin | BGC0001571 | 7 |

## 4- Discussion

Comparing quality metrics such as the number of contigs, N50, and N75 between the assembled genome in this study and the reference genome of *T. trachyspermus* strain 4041, reveals differences in size and quality. The assembly size obtained is 31.3 Mb, smaller than the reference genome size of 32 Mb. The reference genome was sequenced with Oxford nanopore technology, and the genome coverage was 57.0X, whilst Illumina platform with coverage of 20 was used in this study. The reference genome consists of 14 contigs and does not include assembled chromosomes. Its contig N50 is 3.8 Mb. The same determinants for our assembly were 385,981 and 552, respectively. The quality difference between the two genome assemblies may be attributed to variations in sequencing platforms, sequencing depth, and assembly techniques. Compared to long-read techniques such as Oxford Nanopore or PacBio, short-read-based technologies such as Illumina produce genome assemblies with smaller values of N50 and higher counts of contigs (Rayamajhi et al., 2022). Also, low depth or coverage of reads may result in some gaps in the assembly; consequently, some parts of the genome may not be represented in the genomic data (Baker, 2012). This problem occurs particularly in genomic regions with high GC content and regions with high repeats (Ross et al., 2013).

Among SM-BGCs identified, Polyketide synthases (T1PKS) and Nonribosomal peptide synthetases (NRPS / NRPS-like) had the most frequency. Polyketides have a large variety and several polyketides have been reported to be produced by *Talaromyces* species. Among them, a new derivative of spiculisporic acid, spiculisporic acid E, was isolated from the culture of *T. trachyspermus* strain KUFA 0021(Kumla et al., 2014). Polyketides production has been reported by other *Talaromyces* species, including *T. lutcus, T. luteus, Talaromyces* sp., *T. wortmanii, T*.*ardifaciens, T. helices* and *T. flavus (*Zhai et al., 2016).

The production of two cyclic peptides, talaromins A and B, has been reported from the endophytic fungus *T. wortmannii* (Zhai et al., 2016). Different secondary metabolites were detected in the genome, though the similarity percentage ranged between 7% and 100%. A similar FungiSmash analysis was conducted for the genome assembly of *Talaromyces* sp. DC2 revealed the existence of Choline, YWA1, and Squalestatin S1 in the genome of that isolate wich in common with our results. They identified 20 SM-BGCs (Quan et al., 2024). Genome assembly of *Talaromyces pinophilus* strain 1–95 revealed 68 secondary metabolism gene clusters, mainly belonging to T1 polyketide synthase genes and nonribosomal peptide synthase genes (Li et al., 2017). In another research, 62 gene clusters (724 genes) involved in the secondary metabolism, including 18 T1PKS (type 1 PKS), 15 NRPS-like, 9 NRPS, 9 terpenes, 4 NRPS-T1PKS, 4 NRPS-like-T1PKS, 1 *β*-lactone, 1 NRPS-*β*-lactone, and 1 other, were identified in the genome of *T. albobiverticillius*. Among their identified secondary metabolites, six PKS genes were found with 100% similarity in the T1PKS gene cluster. YWA1 was in common with our study (Wang et al., 2023). Genome analysis of *T. verruculosus* SJ9 showed that this genome contained 19 clusters in Type I polyketide synthase (T1PKS), three clusters encoding non-ribosomal peptide synthase cluster (NRPS), 8 clusters in terpene and 13 clusters in NRPS-like (Fu et al., 2024). The nidulanin A gene cluster is conserved in *Aspergillus* and *Penicillium* species (Gonçalves et al., 2021). This compound has yet to be tested for antimicrobial or virulence-related properties (Raffa and Keller, 2019). Fusarins are mycotoxins produced by the genus Fusarium. We have found genes with the potential to code for this polyketide. The similarity of the sequences analysed by FungiSmash for Fusarin is 100%. No previous report of Fusarin encoding genes in Talaromyces or Penicillium species exists. Therefore, this result demands further studies to investigate the real existence and/or expression of the gene encoding this mycotoxin in the genome of the Talaromyces isolate we studied. YWA1 is a fungal monomer Naphtho-γ-pyrone (NGP). It has been demonstrated that NGPs have different biological effects, including anti-viral, anti-microbial, insecticidal, and anti-estrogenic activities (Choque et al., 2015). Dimethylcoprogen is a non-ribosomal peptide siderophore that facilitates the utilisation of iron by fungi in the environment. This siderophore has been reported to be a chelator with a high affinity towards iron. This siderophore has been shown to inhibit the growth of several bacterial and fungal species (Renshaw et al., 2002).

Two compounds belonging to the terpenes were identified in the *T. trachyspermus* genome: squalestatin S1 (SQS1) and demethoxyviridin. SQS1, or zaragozic acid (ZA), is a fungal metabolite with broad antifungal activity. It is also a lead compound for cholesterol-lowering drugs (Lebe, 2020). It has been demonstrated that demethoxyviridin and some synthetic analogs, such as 1α-Hydroxydemethoxyviridin, have antibacterial and antifungal activities (Kiran et al., 2005).

Using FungiSmash analysis, we identified sequences similar to Phyllostictine A/ Phyllostictine B (a similarity percentage of 40% and 30 %). Phyllostictine A is a powerful toxin produced by the fungus *Phyllosticta* cirsii, a potential natural mycoherbicide of *Cirsium* arvense or Creeping thistle as an important weed in the world (Zonno et al., 2008). In 2008, Evidente and coworkers reported four new classes of natural herbicides named phyllostictines A-D, produced by the fungus *Ph. cirsii*; among them, phyllostictine A was demonstrated to be the most potent member, showing very high efficacy against thistles. They also showed that phyllostictine A is more efficient than fusaric acid, a powerful toxin, and faster acting than glyphosate (Evidente et al., 2008). Another study in 2011 discovered the anti-cancer activity of phyllostictine A against several human tumour cell lines (Le Calvé et al., 2011).

Scytalidin is another antifungal secondary metabolite detected with 18 percent similarity in the genome of our studied isolate. This fungitoxic compound is produced with Scytalidium species (Strunz et al., 1972). The low similarity of the sequences could be due to the assembly quality and/or different compounds with similar sequences.

FungiSmash also determined a sequence with 14 percent similarity to another herbicidal compound coexisting. This metabolite has been purified from the cultures of *Paecilomyces variotii* SANK 21086 in 1991. Based on its herbicidal features, it was classified as a postemergence herbicide against certain young annual and perennial monocotyledonous and dicotyledonous weeds with no adverse effect on corn seedlings (Nakajima et al., 1991).

There is also one detected sequence in the studied genome, similar to elsinochrome C, a light-active non-host selective phytotoxin compound produced by the plant pathogenic fungus *Elsinoë arachidis* (Jiao et al., 2019). No phytotoxicity has been reported in the previous study against the tomato plants as the host of broomrape in greenhouse experiments of inoculating the pots with the studied isolate of *T. trachyspermus* IRAN 3054C to control broomrape by Hemmati and Gholizadeh (2019). Also, in another greenhouse experiment to investigate the biocontrol effects of this isolate against *S. sclerotiorum* on tomato, we did not observe any phytotoxicity against the host plant (unpublished data). We have also conducted a pathogenicity test of the mentioned isolate on a few plants and have not observed any pathogenicity or phytoxicity. The similarity between sequences was only 12 %, which is not a supportive percentage. Whereas the similarity percentages of the query sequences of the studied genome with the detected secondary metabolites from FungiSmash analysis are variable between 7% and 100%, it is necessary to conduct a metabolomics analysis for all determined metabolites of this study.

The studied isolate’s genome contains a high number of fungal cell wall-degrading enzymes, including chitinases, glucanases, peptidases, proteases, and lipases, which are essential for antagonistic activity. This isolate also contains a high number of phosphatase coding sequences and five domains for siderophore synthase proteins in its genome. Phosphate insolubilisation and siderophore production are significant mechanisms of plant growth promotion. Previous studies have reported the production of these enzymes and phosphate insolubilization by *T. trachyspermus* (Sahu et al., 2019). Also, this isolate has been reported to produce high amounts of siderophores (Sahu and Prakash, 2021). Plant growth promotion activity has also been reported by another *Talaromyces* biocontrol species, *T. pinophilus*, which has promoted rice growth (Khalmuratova et al., 2015). Genome analysis of several other species of *Talaromyces* has also revealed that *Talaromyces* species have gene clusters associated with cell wall-degrading enzymes. Whole genome sequencing of *T. piceus* strain 9-3 showed that its genome had different lignocellulolytic enzymes, including two cellobiohydrolases, ten β-glucosidase, and one endo-β-1,4-glucanase gene cluster (He et al., 2017). *T. pinophilus* strain 1–95 contained a high number of CAZYmes in its genome, including eight β-1,4-endoglucanases, two cellobiohydrolases, 29 β-glucosidases, 97 hemicellulose-degrading enzymes, and 24 α-amylases (Li et al., 2017). The high number of glucosidase and glucanases in the genome of other *Talaromyces* species is consistent with our findings. *Talaromyces* sp. strain DC2 contained 653 CAZymes responsible for plant cell wall degradation; among them, β-glucosidases and β-galactosidases had 27 genes and 23 genes in the genome. The most abundant glycoside hydrolases (GH) belonged to chitinases (Quan et al., 2024). Study on genome assembly of *T. albobiverticillius* revealed that glycosed hydrolases including chitinases (GH18), *β*-glucosidases (GH3) and polygalacturonases (GH28), were the most frequent CAZYmes in its genome (Wang et al., 2023). We also report high copies of chitinases in our studied genome. Studies on CAZYmes of most Talaromyces species have mainly focused on plant cell wall degradation, as they have studied the isolates for plant debris degradation potential, whereas, in current research, we searched our studied genome for enzymes effective on fungal cell wall hydrolysing, plant cell wall degradation, and mycoparasitism. A chromosome-level genome assembly of *Talaromyces rugulosus* was generated using a combination of PacBio long-read and Illumina paired-end data. *T. rugulosus* is a powerful enzyme producer and also a promising biocontrol agent against *Aspergillus flavus*, a notorious mycotoxin producer plant pathogen (Wang et al., 2020). Their results showed that the genome of *T. rugulosus* is rich in genes encoding proteases, carbohydrate-active enzymes, fungal cell wall–degrading enzymes, and secondary metabolite biosynthetic genes, demostrating its mycoparasitic capability (Wang et al., 2020).

## Conclusion

The genome of *T. trachyspermus*, isolate IRAN 3054C, is rich in secondary metabolite gene clusters, encoding genes for proteases and fungal cell wall degrading enzymes, reflecting its mycoparasitic potential against plant pathogenic fungi. Its genome contains sequences similar to genes encoding antifungal and herbicidal secondary metabolites, which is consistent with the in vitro and greenhouse observations reported for this isolate against *O*.*ramosa* (Hemmati and Gholizadeh, 2019 and unpublished data). YWA1 and Dimethylcoprogen, as siderophores with antifungal and antibacterial activity, were detected in the genome with 100 % similarity. Other antifungal or antimicrobial SMs with a similarity of < 100% are squalestatin S1 (60%), nidulanin A (50%), scytalidin (18%), and demethoxyviridin (7%). Therefore, for the last groups of SMs, it is necessary to conduct a metabolomics study to find out the exact components produced with this isolate. Phyllostictin A/B (30 and 40 %) and Cornexistine (14%) are herbicidal SMs. According to the low similarity percentage, their production also needs to be investigated in a study of the metabolites of this isolate. Genome analysis showed that strain IRAN 3054C serves as a potential source of essential enzymes for antagonistic activity against fungal pathogens. Although its genome harbours other CAZYmes for plant cell wall degradation, the number of genes encoding for mycoparasitism in the genome of this starin is prevalent. According to the genomic data, we can conclude that *T. trachyspermus* IRAN 3054C employed antibiosis and parasitism against fungal pathogens. However, its mechanism for biocontrol against broomrape (*O*.*ramosa*) is still unknown, and further investigations are required to confirm the distinct mechanisms of its herbicidal effect.

## Declarations

### Ethics approval and consent to participate

Not applicable

### Consent for publication

Not applicable

### Authors’ contributions

R. H. wrote the main manuscript and conducted data analysis and part of the technical lab work; A. D. made major edits in the text of the manuscript and has also helped in some parts of lab work; S. S. helped with technical laboratory work in Iran; J.B. has provided laboratory and scientific facilities, and supported sequencing costs. All authors reviewed the manuscript.

### Data availability

Sequence data that support the findings of this study have been deposited in DDBJ/ENA/GenBank under the accession JBNAEF000000000.

## Acknowledgements

Not applicable

